## Supplementary Material for "Simulating neural network criticality and resource dynamics with Rydberg gases"

### Supplementary Information for Simulating neural network criticality and resource dynamics with Rydberg gases

#### CALCULATING THE BRANCHING RATIO

The goal of this section is to calculate the average number of facilitated excitations $BR$ that a seed excitation will create in the mean-field picture. For that, we assume that the seed excitation decays with rate $\gamma$ and the dynamics of a second test atom initially in the ground state and at distance $r$ from the seed atom can be described by the optical Bloch equations.

We're interested in the state of the second atom at the moment when the seed decays. We start with the generic solution of the optical Bloch equations for an atom initially in the ground state as given in equation (10) of [1]:

$$w = Ae^{-at} + \left[B\cos(qt) + \frac{C}{q}\sin(qt)\right]e^{-pt} + w_s$$

Where $w$ is the imbalance $w = \rho_{gg} - \rho_{ee}$. Here $w_s = \frac{\Delta^2+\gamma^{*2}}{\Delta^2+\gamma^*(\gamma^*+\Omega^2/\gamma)}$ is the steady-state imbalance given in equation (3) of [1] and $A$,$B$,$C$,$a$,$p$ and $q$ are prefactors depending on Rabi frequency $\Omega$, local detuning $\Delta(r)$, decay rate $\gamma$ and dephasing $\gamma^*$, and are explicitly given in equation (11) of [1]:

$$A = \frac{1}{D}\left[-\Omega^2 + (p^2+q^2)(1-w_s)\right]$$
$$B = \frac{1}{D}\left[a(a-2p)(1-w_s) + \Omega^2\right]$$
$$C = \frac{1}{D}\left[-\Omega^2(a-p) + a(ap-p^2+q^2)(1-w_s)\right]$$
$$D = (a-p)^2 + q^2$$

And equation (6) of [1] gives the values for $a$, $p$ and $q$:

$$a = \alpha - (S_+ + S_-),$$
$$p = \alpha + \frac{1}{2}(S_+ + S_-),$$
$$q = \frac{\sqrt{3}}{2}(S_+ - S_-)$$
$$\alpha = \frac{2\gamma^* + \gamma}{3},$$
$$S_\pm = \left(R \pm \sqrt{R^2+Q^3}\right)^{1/3}$$

Our seed atom decays at time $t$ with probability $\gamma e^{-\gamma t}$. The probability to find our test atom in the excited state once this happens can be written as $\frac{1-w}{2}$. To obtain the probability $P_{\text{fac}}(\Omega, \gamma, \gamma*, \Delta(r))$ that the test atom gets facilitated, we integrate over all possible decay times $t$:

$$\begin{aligned}
P_{\text{fac}} &= \int_0^\infty \frac{1-w}{2}\gamma e^{-\gamma t}dt \\
&= \frac{\gamma}{2}\int_0^\infty \left(1 - Ae^{-at} - \left[B\cos(qt) + \frac{C}{q}\sin(qt)\right]e^{-pt} - w_s\right)e^{-\gamma t}dt \\
&= \frac{\gamma}{2}\left((1-w_s)\int_0^\infty e^{-\gamma t}dt - A\int_0^\infty e^{-(\gamma+a)t}dt - B\int_0^\infty \cos(qt)e^{-(\gamma+p)t}dt - \frac{C}{q}\int_0^\infty \sin(qt)e^{-(\gamma+p)t}dt\right) \\
&= \frac{\gamma}{2}\left((1-w_s)\frac{1}{\gamma} - \frac{A}{\gamma+a} - \frac{B(\gamma+p)}{(\gamma+p)^2+q^2} - \frac{Cq}{q((\gamma+p)^2+q^2))}\right)
\end{aligned}$$

Our derivation includes off-resonant excitation, which is always present regardless of any seed atoms. We therefore need to subtract this off-resonant rate in order to obtain the effect of our seed atom. The total branching ratio $BR$ can be derived by integrating this probability over the volume of the system and the density $n$ of ground state atoms. The local detuning $\Delta(r)$ is given by the laser detuning $\Delta$ and the $c_6$ interaction as $\Delta(r) = \Delta - \frac{C_6}{r^6}$:

$$BR = \int_{R^3} n(R) \left[P_{\text{fac}}\left(\Omega, \gamma, \gamma*, \Delta - \frac{C_6}{r^6}\right) - P_{\text{fac}}\left(\Omega, \gamma, \gamma*, \Delta\right)\right] dV$$
$$= 4\pi n \int_0^\infty r^2 \left[P_{\text{fac}}\left(\Omega, \gamma, \gamma*, \Delta - \frac{C_6}{r^6}\right) - P_{\text{fac}}\left(\Omega, \gamma, \gamma*, \Delta\right)\right] dr$$

Setting $BR = 1$ allows solving for the critical density $n_c$:

$$n_c = \frac{1}{4\pi \int_0^\infty r^2 \left[P_{\text{fac}}\left(\Omega, \gamma, \gamma*, \Delta - \frac{C_6}{r^6}\right) - P_{\text{fac}}\left(\Omega, \gamma, \gamma*, \Delta\right)\right] dr} \quad (1)$$

A numerical solution of this integral is shown as yellow line in Fig. 2.

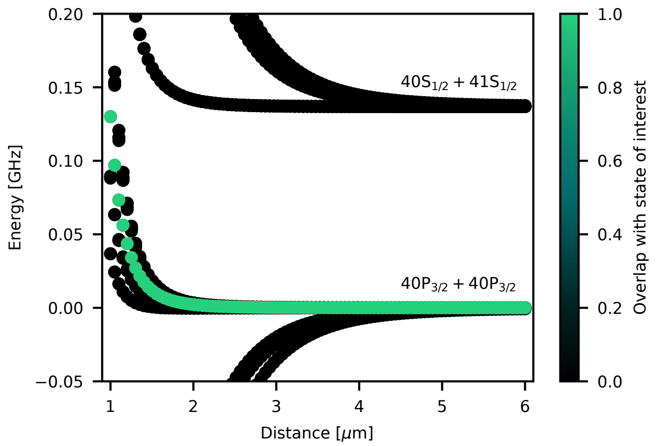


Supplementary Figure 1. Rydberg interaction potential. Calculation is done using the pairinteraction software package [2]. The different 40P curves correspondent to different $m_j$ quantum numbers. Which one our coupling laser excites depends on the angle between inter-atomic axis (which defines the quantization axis, as the Rydberg-Rydberg interaction is our largest energy scale) and laser polarization. The 40S+41S pair state at +137 MHz detuning causes additional features in the detuning dependent phase diagram shown in Fig. 2.

#### INTERACTION POTENTIAL

We calculate the Rydberg-Rydberg interaction using the pairinteraction [2] software package. The potential curves are shown in Supplementary Fig. 1. Fitting a $C_6/r^6$ potential to the resulting curve allows us to obtain a value of $C_6 = 0.111\,\text{GHz}\,\mu\text{m}^6$ and a facilitation distance of $r_{\text{fac}} = 1.185\,\mu\text{m}$ at a laser detuning of $\Delta = 40\,\text{MHz}$.

#### SUPPLEMENTARY REFERENCES


*



[1] H.-R. Noh and W. Jhe, Analytic solutions of the optical Bloch equations, Optics Communications **283**, 2353 (2010).

[2] S. Weber, C. Tresp, H. Menke, A. Urvoy, O. Firstenberg, H. P. Büchler, and S. Hofferberth, Calculation of Rydberg interaction potentials, Journal of Physics B: Atomic, Molecular and Optical Physics **50**, 133001 (2017).
